# Single-cell resolution analysis of the crosstalk between chemogenically activated astrocytes and microglia

**DOI:** 10.1101/2020.04.27.064881

**Authors:** Stéphanie Philtjens, Marion T. Turnbull, Brian P. Thedy, Younghye Moon, Jungsu Kim

**Affiliations:** Department of Medical and Molecular Genetics, Indiana University School of Medicine, Indianapolis, IN 46202, USA; Stark Neuroscience Research Institute, Indiana University School of Medicine, Indianapolis, IN 46202, USA; Department of Neurology, Mayo Clinic, Jacksonville, FL 32224, USA; Department of Neuroscience, Mayo Clinic, Jacksonville, FL 32224, USA

**Keywords:** single cell RNA sequencing, DREADDs, astrocytes, microglia, chemogenetics, hippocampus, cortex

## Abstract

Astrocytes are the most common glial cell type in the brain, yet, it is unclear how their activation affects the transcriptome of neighboring cells. Engineered G protein-coupled receptors (GPCRs) called Designer Receptors Exclusively Activated by Designer Drugs (DREADDs) enable selective activation of specific cell types, such as astrocytes. Here, we combine activation of astrocytes in the hippocampus and cortex of healthy mice with single-cell RNA sequencing. Our data show that long-term activation of astrocytes dramatically alters the transcriptome of astrocytes and microglia. Genes that were differentially expressed in Gq-DREADD-activated astrocytes are involved in neurogenesis and low-density lipoprotein particle biology, while those in the microglia were involved in lipoprotein handling, purinergic receptor activity, and immune cell migration and chemotaxis. Furthermore, network analysis showed that Gq-DREADD-mediated activation in astrocytes resulted in an upregulation of genes involved in the GPCR signaling pathways and calcium ion homeostasis, confirming astrocyte activation. This dataset will serve as a resource for the broader neuroscience community, and our findings highlight the importance of studying transcriptomic alterations in microglia after astrocyte activation *in vivo*.

## Introduction

Astrocytes are the most abundant glial cell in the brain and affect most aspects of neural function. They are responsible for neurotransmitter and ion homeostasis at the level of the synapse, provide trophic support to neurons, regulate blood flow, and provide energy substrates (Abbott et al., 2006; Sofroniew and Vinters, 2010). Correspondingly, in disease states, astrocytes can undergo morphofunctional remodeling called ‘reactive astrocytosis’, where normal homeostatic mechanisms are lost and pro-inflammatory responses are upregulated (Liddelow and Barres, 2017; Ridet et al., 1997; Zamanian et al., 2012). This reactive phenotype may contribute to disease progression or may even trigger initial pathological changes such as tauopathy and neuronal cell death, making reactive astrocytes an important cell type to study in neurodegenerative diseases such as Alzheimer’s disease (AD) (Chun et al., 2020; Richetin et al., 2020; Sofroniew and Vinters, 2010). Thus, a better understanding of how changes in astrocyte function affects surrounding cells *in vivo* would be critical for our understanding of normal physiology and disease progression.

To fully characterize the complex molecular interactions between astrocytes and neighboring cells *in vivo*, we used a combination of chemogenetics and single-cell RNA sequencing (scRNA seq). Expression of engineered G protein-coupled receptors (GPCRs), called Designer Receptors Exclusively Activated by Designer Drugs (DREADDs), in astrocytes allows for activation of this cell type through selective binding of its ligand (Lee et al., 2014; Roth, 2016; Urban and Roth, 2015). These mutated muscarinic receptors respond exclusively to clozapine-N-oxide (CNO) and are unresponsive to their endogenous ligand acetylcholine (Armbruster et al., 2007; Rogan and Roth, 2011). In this study we used the Gq-coupled human muscarinic type 3 receptor (hM3Dq) whose activation results in Ca^2+^ mobilization and ERK1/2 phosphorylation (Armbruster et al., 2007). The *in vivo* functionality of DREADD-mediated astrocyte activation has previously been demonstrated in mice, with excitation of hM3D-expressing astrocytes in the CA1 region of the hippocampus resulting in increased neuronal activity and enhanced memory (Adamsky et al., 2018). However, the effect of astrocyte activation on the transcriptome of neighboring cells was not investigated.

Recently, the increased interest in astrocytes in health and disease, as well as the progress in scRNA seq technologies resulted in an increased knowledge of the transcriptional complexity of astrocytes (Batiuk et al., 2020; Bayraktar et al., 2020; Das et al., 2020; Habib et al., 2020; Pan et al., 2020). The majority of studies looked at the transcriptional profile of astrocytes in disease models and have shown transcriptional differences between astrocytes associated with Huntington’s disease (Al-Dalahmah et al., 2020; Diaz-Castro et al., 2019), AD (Habib et al., 2020; Xu et al., 2020), or aging (Pan et al., 2020). They showed that, although there were commonalities between the different transcriptional phenotypes, there were also important differences in the number and type of genes that were transcriptionally altered (Sofroniew, 2020). In addition, while most of these studies focused on ‘reactive’ astrocytes, a recent scRNA seq study looked at ‘normal’ astrocytes in C57BL/6J mice at post-natal day 56. They showed the presence of transcriptionally distinct astrocytes in the cortex and hippocampus, and showed the presence of spatial effects on the transcriptome of these astrocytes (Batiuk et al., 2020).

In this study, we were interested in the transcriptional changes that occur in astrocytes and neighboring cells after astrocyte activation. To selectively activate astrocytes, we used adeno-associated virus transduction of Gq-DREADD into the hippocampus and cortex of C57BL/6J mice under the control of the glial fibrillary acidic protein (GFAP) promoter. CNO was administered over an eight-week period to simulate chronic activation of astrocytes instead of initiating an acute injury response. We profiled a total of 34,736 cells and showed that Gq-DREADD-mediated activation in astrocytes does not only change the transcriptional profile of the astrocytes themselves, but also of neighboring microglia, as can be expected. To the best of our knowledge, our study is the first to investigate the effect of Gq-DREADD activated astrocytes on surrounding brain cells at the single cell level using scRNA seq.

## Material and Methods

### Plasmid preparation and AAV packaging

pAAV-GFAP-hM3D(Gq)-mCherry was obtained from Addgene (Construct: Bryan Roth; Addgene, plasmid # 50478; http://n2t.net/addgene:50478; RRID:Addgene_50478). Plasmid DNA was prepared and purified using a QIAGEN Plasmid Maxi Kit (Qiagen, Cat. No. 12163) per the manufacturer’s protocol. AAV particles were packaged into serotype 5 type capsid and purified using standard methods (Zolotukhin et al., 1999). Briefly, AAV was generated by co-transfection with the cis plasmid pDP5 (Plasmid Factory, AAV5 helper plasmid) into HEK293T cells. Cells were harvested 72 h after transfection, treated with 50 Units/mL Benzonase (Sigma Aldrich), and lysed by freeze thaw. The virus was then purified using a discontinuous iodixanol gradient and buffer exchanged to phosphate buffered saline (PBS) using an Amicon Ultra 100 Centrifugation device (Millipore). The genomic titer of each virus was determined by quantitative PCR using the ABI 7700 (Applied Biosystems) and primers specific to the WPRE. The viral DNA samples were prepared for quantification by treating the virus with DNaseI (Invitrogen) and Proteinase K, and samples were compared against a standard curve of supercoiled plasmid diluted to 1^4^ to 1^7^ copies per mL. AAV5 is astrocyte specific and its injection into the striatum and substantia nigra has previously demonstrated astrocyte-specific expression with no expression in neurons or microglia (Drinkut et al., 2012; Merienne et al., 2013).

### Animals

All experiments were approved by the Mayo Clinic Institutional Animal Care and Use Committee (IACUC). Male C57BL/6J mice (n=4, The Jackson Laboratory, stock number #000664) were group housed and allowed to acclimate to their new environment for 1 week before surgery. Mice were housed on a 12-hour dark/light cycle with water and standard chow available *ad libitum*.

### Stereotaxic Surgery

Eight-week-old mice (19-24 g, n = 2/group) were anesthetized by intraperitoneal (i.p.) injection of ketamine (100 mg/kg) and xylazine (10 mg/kg) and placed on a stereotaxic instrument. The skin was opened and a hole in the skull was made by a handheld drill. The needle was lowered into the hippocampus (Anterior-Posterior (AP): -2 mm, Medial-Lateral (ML): ±1.5 mm, Dorsal-Ventral (DV): -1.7 mm from the Bregma) and the virus (AAV-GFAP-hM3D(Gq)-mCherry; Serotype 5; 1.0×10^12^ genome copies/mL) was infused at a rate of 0.4 μL/min (1.5 μL total volume). The needle was allowed to remain in place for 5 minutes then slowly raised 0.9 mm to the cortex (AP: -2 mm, ML: ±1.5 mm, DV: -0.8 mm from the Bregma) and more virus was infused (0.4 μL/min; 1.0 μL total volume). The needle was again left for 5 minutes before being removed. This was repeated on the contralateral hemisphere for bilateral injection of the virus into both hippocampi and cortices (Figure 1A).

**Figure 1:**
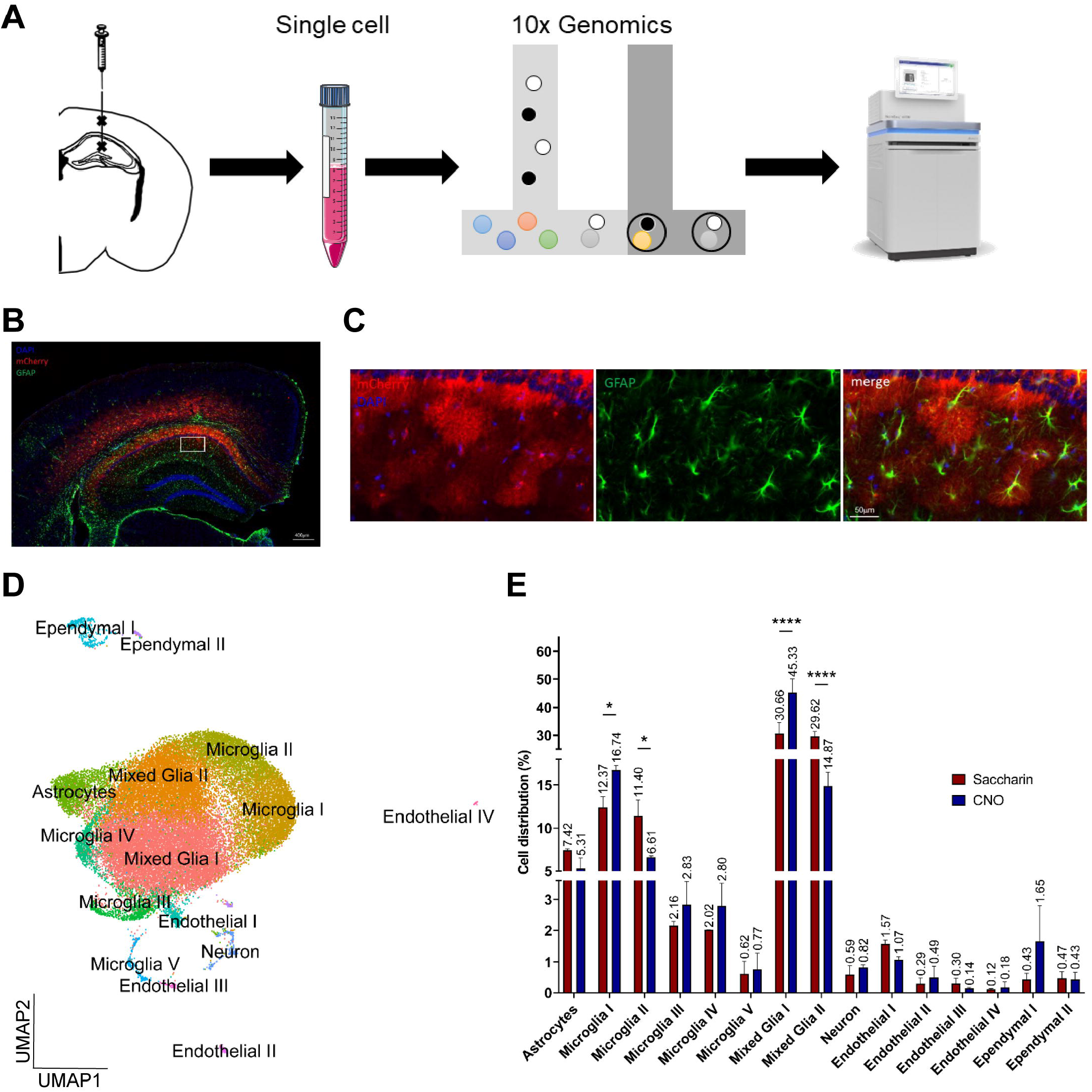
scRNA sequencing shows differences in cell distribution in the hippocampus and cortex of Gq-DREADD-injected mice. **(A)** Schematic presentation of the scRNA seq workflow. Male C57BL/6J mice were transduced with AAV5-GFAP-hM3D(Gq) at 8 weeks of age and treated with saccharin or CNO in drinking water for 8 weeks. **(B)** Representative fluorescence image of the hippocampus and cortex demonstrating Gq-DREADD virus spread (red; mCherry) and astrocytes (green; GFAP). Cell nuclei are indicated by blue fluorescence (DAPI). **(C)** Magnified view (from white box in **(B)**) demonstrating mCherry specificity to astrocytes. **(D)** UMAP plot showing the detected cell types in the anterior hippocampus and anterior cortex. Cluster names were assigned using conserved marker genes. **(E)** The cell distribution of the detected cell clusters in the saccharin and CNO group. Data represent mean ± standard deviation, two-way ANOVA and Bonferroni’s multiple comparisons test. * *p* ≤ 0.05; **** *p* ≤ 0.0001

### Chronic CNO Administration

CNO (Hello Bio, Cat. No. HB1807) was dissolved in clean drinking water at 0.25 mg/mL to approximate a 5mg/kg/day dose of CNO. 5 mM of saccharin (Sigma Aldrich, Cat. No. 109185) was added to mask the bitter taste of CNO and control groups had only saccharin in their drinking water. Mouse cage water containing CNO and saccharin was carefully monitored and changed for fresh CNO/saccharin every two to three days. CNO/saccharin was given for 8 weeks after stereotaxic surgery.

### Tissue Preparation and Immunohistochemistry

For tissue that was to be assessed histologically, mice were sacrificed by ketamine (100mg/kg i.p.) overdose followed by transcardial perfusion with 4% paraformaldehyde (PFA). Tissue was post-fixed in 4% PFA overnight at 4°C, and then left for 24 h in PBS containing 30% sucrose. Tissue was then embedded in OCT tissue freezing medium (Tissue-Tek®, Cat. No. 25608-930) and sections were cut in the coronal plane (30 μm) using a cryostat.

Fluorescence immunohistochemistry was performed on free floating coronal sections to identify mCherry-positive cells and GFAP-positive astrocytes, NeuN-positive neurons or Iba1-positive microglia in the hippocampus and cortex. Briefly, sections were blocked for 1 h at room temperature in 5% bovine serum albumin (BSA), and 0.1% Triton X-100 in PBS followed by incubation overnight in rabbit anti-GFAP antibody (1:1000; Thermo Fisher Scientific, Cat. No. 2.2B10, Figure 1B and 1C), mouse anti-NeuN (1:1000; Novus, Cat. No. NBP1-92693, Supplementary Figure 1A) or rabbit anti-Iba1 (1:1000; Wako, Cat. No. 019-19741, Supplementary Figure 1B), and rabbit anti-mCherry (1:1000; Abcam, Cat. No. ab167453). After washing in 0.1% Triton X-100 in PBS, sections were incubated in AlexaFluor 488, 594 and 647 (1:1000; Invitrogen) and DAPI (1:1000; Thermo Scientific) for 2 h. Sections were then washed and coverslipped with Prolong Antifade Diamond Mountant (Invitrogen). Fluorescence microscopy was performed using a fluorescence slide scanner (Aperio, Leica Biosystems).

### Isolation of cells

Mice were sacrificed by ketamine (90 mg/kg i.p.) and xylazine (10 mg/kg) overdose followed by transcardial perfusion with ice-cold PBS (approx. 20 mL). Hippocampal and cortical tissue containing the virus was dissected out from the freshly isolated brain. The tissue was minced with a razor blade and incubated for 30 min in ice-cold accutase (StemPro Accutase, Fisher Scientific, Cat. No. A1110501) at 4°C. The dissociated tissue was centrifuged at 300*g* for 10 min at 4°C. The pellet was resuspended in ice-cold Hanks’ Balanced Salt solution (Life Technologies, Cat. No. 14025-092) and triturated with a 15 mL serological pipette. This step was repeated until the entire tissue was dissociated. Next, the sample was filtered first through a 70 μm cell strainer, followed by a 40 μm cell strainer. The cells were centrifuged at 300*g* for 5 min at 4°C.

### Myelin removal

Myelin removal was performed following the Miltenyi’s Myelin Depletion Protocol. In short, the pellet obtained during cell isolation was resuspended in magnetic-activated cell sorting (MACS) buffer (0.5% BSA in PBS, Thermofisher, Cat. No. AM2618), myelin removal beads (Miltenyi Biotec, Cat. No. 130-096-733) were added, and the solution was incubated for 15 min at 4°C. The cells were centrifuged at 300*g* for 10 min at 4°C, the pellet was resuspended in MACS buffer and the solution was passed through LS columns (Miltenyi Biotec, Cat. No. 130-042-401). The cells were centrifuged at 300*g* for 15 min at 4°C, the pellet was resuspended in MACS buffer, centrifuged again at 300*g* for 15 min at 4°C and resuspended in PBS containing 0.01% BSA.

### Single cell RNA sequencing

The obtained single cells were subjected to droplet-based single cell RNA sequencing. The 10x Genomics Chromium™ Single Cell 3’ Library & Gel Bead Kit v2 was used per manufacturer’s instructions. The libraries were sequenced on an Illumina NovaSeq 6000 SP at the Indiana University Center of Medical Genomics (Figure 1A). The 10x Genomics software Cell Ranger v.3.0.2 was used for sample demultiplexing, barcode processing and single cell counting. All samples were aligned against the mouse reference genome (mm10).

Subsequent data analysis was performed using the R (v.3.5.2) package Seurat v.3.0.2 (Butler et al., 2018; Stuart et al., 2019). Standard quality control included the removal of cells with a mitochondrial count >5%, as well as cells with less than 200 and more than 2,000 detected genes (Supplementary Figure 1C). A total of 34,736 cells remained after quality control filtering steps. The data were log-normalized using a scaling factor of 10,000, scaled and regressed against the percentage of mitochondrial content. Principal component analysis (PCA) was performed using the top 3,000 most variable genes and the number of PCAs to use was determined after 1,000 permutations using an ElbowPlot. The first 21 PCAs were used to determine the K-nearest neighbor (KNN) and cluster the cells into 15 different cell clusters. The clustering resolution used was 0.5. Marker genes for naming the clusters were determined using the FindConservedMarkers function in Seurat (Supplementary Data 1). Cells from the glial clusters (astrocytes, microglia I-V, mixed glia I, mixed glia II) were taken for re-clustering.

### Analysis of differential gene expression

Differential gene expression analysis between the saccharin group and the CNO group was done in Seurat using a likelihood test assuming an underlying negative binomial distribution. Genes with a p-value < 0.05 after Bonferroni correction were considered to be significantly differentially expressed between both groups. The differentially expressed genes were analyzed for functional enrichment using g:Profiler (accessed on July 2-4, 2020, version e99_eg46_p14_f929183) (Raudvere et al., 2019), and network analysis was performed using the Analyze Networks algorithm in MetaCore™ (Clarivate Analytics, v.19.4 build 69900).

### Statistical analysis

Two-way ANOVA and Bonferroni’s multiple comparisons test was performed using GraphPad Prism version 9.0.0 for Windows (GraphPad Software, San Diego, California USA, http://www.graphpad.com). Data was represented as mean ± standard deviation. * *p* ≤ 0.05; **** *p* ≤ 0.0001

### Data availability

Full list of differentially expressed genes and the g:Profiler analysis are provided online as Supplementary Data. The raw sequencing and processed data are available through the GEO database (GSE154208). This data will be made available to the public upon publication of this manuscript. However, the data can be made available to the reviewers for the period of the reviewing process.

## Results

### Astrocyte-Specific Expression of Gq-DREADD

Microinjection of AAV5-GFAP-hM3D(Gq) resulted in virus spread around the hippocampal and cortical injection points (Figure 1B). Histological visualization with the fluorescent reporter mCherry confirmed that Gq-DREADD expression remained restricted to astrocytes based on their co-localization with the astrocyte marker GFAP (Figure 1C) and morphology consistent with astrocytic immunoreactivity patterns (Benediktsson et al., 2005; Bushong et al., 2002; Scofield et al., 2015). Moreover, mCherry did not show co-localization with the neuronal marker, NeuN, or the microglial marker, Iba1 (Supplementary Figure 1A and 1B).

### Single cell RNA sequencing identifies 15 transcriptionally different cell types

To determine how activation of astrocytes affects the transcriptome of astrocytes and neighboring microglia, we performed scRNA seq on cells isolated from the hippocampus and cortex of four-month-old C57BL/6J mice. These mice were injected with AAV5-GFAP-hM3D(Gq) at 8 weeks of age (Figure 1A) and treated with CNO or saccharin for 8 weeks. Two samples per condition were sequenced with an average of 442M reads per sample, or an average of 37,000 reads per cell. About 56% of all reads were exonic, while 29% were intronic and only 5% were intergenic. A total of 34,736 cells passed the quality control filters, 16,107 cells in the CNO treated group and 18,629 cells in the saccharin group. Unsupervised clustering resulted in the identification of 15 different cell clusters (Figure 1D, Supplementary Figure 1C and 1D). The cell clusters were annotated manually using marker genes that were conserved between both conditions as astrocytes, microglia, neurons, endothelial cells, ependymal cells, and mixed glia (Supplementary Data 1) (Arneson et al., 2018; McKenzie et al., 2018). We observed an enrichment of glial cells with 96% of all cells detected annotated as either astrocytes, microglia or mixed glia.

When assessing the cell distribution between the CNO treated and the saccharin groups across different cell types, we noticed significant differences in the percentage of cells in the mixed glia I (30.66% in the saccharin group vs. 45.33% in the CNO group, *p* ≤ 0.0001), mixed glia II (29.62% in the saccharin group vs. 14.87% in the CNO group, *p* ≤ 0.0001), microglia I (12.37% in the saccharin group vs. 16.74% in the CNO group, *p* ≤ 0.05) and microglia II clusters (11.40% in the saccharin group vs. 6.61% in the CNO group, *p* ≤ 0.05, Figure 1E, Supplementary Figure 1D and Supplementary Table 1). A smaller difference was observed in the astrocyte cluster (7.42% in the saccharin group vs. 5.31% in the CNO group, Figure 1E, Supplementary Figure 1D and Supplementary Table 1).

### Chronic activation of astrocytes results in a transcriptionally different astrocyte type

Since Gq-DREADD was only expressed in astrocytes, we analyzed the effect of long-term Gq-DREADD activation by comparing the genes that were differentially expressed between the saccharin group and the CNO group in the astrocyte cluster. We identified 396 genes (*p* < 0.05) that were either up- or downregulated in the CNO group compared to the saccharin group (Figure 2A). Gene enrichment and functional annotation analysis showed that transcripts that were upregulated in the CNO group are involved in adenylate cyclase activator activity and the regulation of cell motility and migration, while the downregulated genes were involved in gene expression and translation (Supplementary Data 1). Network analysis was performed using MetaCore™. We found that, among the genes that were upregulated after activation of Gq-DREADD with CNO, 16 of them were in a 50-gene network represented by CMKLR1, SPARCL1, Rich1, Transcobalamin II, OLFML3 (Figure 2B and 2C). The top processes linked to this network were GPCR signaling pathway, ionotropic glutamate receptor signaling pathway, adenylate cyclase inhibiting GPCR signaling pathway, cellular calcium ion homeostasis, and calcium ion homeostasis (Figure 2B). Since these cellular processes are pathways regulated by Gq-DREADD, this network analysis confirms Gq-DREADD-mediated activation of astrocytes in CNO-dosed mice. Figure 2C shows that the genes involved in the network shown in Figure 2B are upregulated in the CNO group compared to the saccharin group. We also compared the transcriptomic profile of our chronically Gq-DREADD activated astrocytes with the transcriptomic profiles of astrocytes in recent publications, as well as with PAN-reactive, and A1 and A2 astrocytes (Supplementary Figure 2A-C) (Das et al., 2020; Habib et al., 2020; Liddelow et al., 2017). However, the transcriptomic profile of Gq-DREADD-activated astrocytes were different from those of PAN reactive, A1, A2, disease-associated astrocytes or any other transcriptomics phenotype studied thus far.

**Figure 2:**
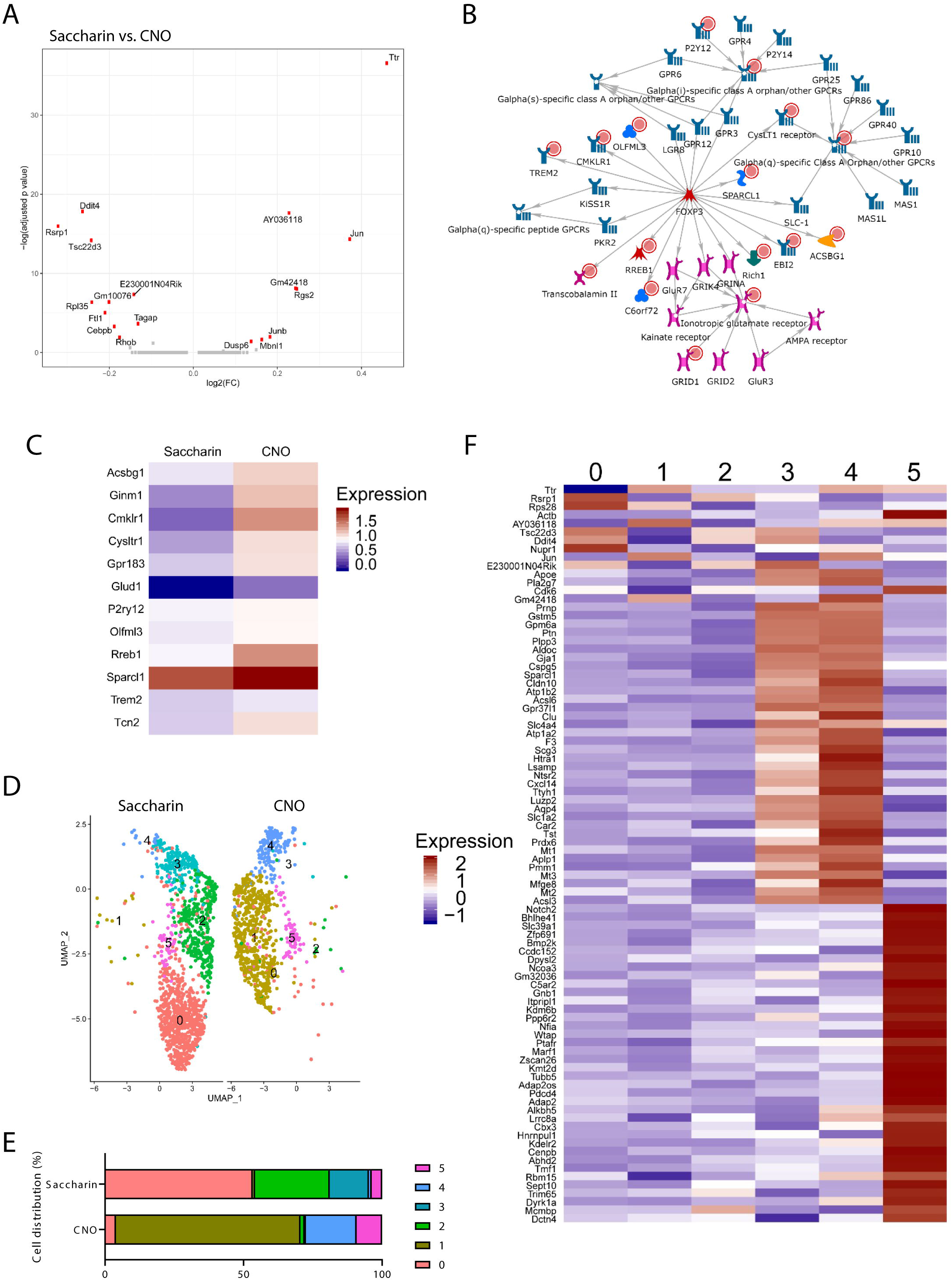
Re-clustering of the astrocyte cell cluster shows the presence of transcriptionally different astrocytes after astrocyte-specific activation with the Gq-DREADD. **(A)** Volcano plot showing differentially expressed genes between the saccharin group and the CNO group in the astrocyte cluster. Differentially expressed genes with an adjusted p value < 0.05 after Bonferroni correction are shown in red. **(B)** Functional network of genes that are upregulated after Gq-DREADD activation in astrocytes. Processes that were enriched are G protein coupled receptor signaling pathway, ionotropic glutamate receptor signaling pathway, adenylate cyclase inhibiting G protein coupled receptor signaling pathway, cellular calcium ion homeostasis, and calcium ion homeostasis. The upregulated genes are marked with red circles. **(C)** Heatmap showing the upregulated expression of genes involved in the network shown in **(B)**. **(D)** UMAP showing the six clusters identified after re-clustering the astrocyte cluster (2,238 cells). The UMAP is split into two groups, the saccharin group (1,380 cells) and the CNO group (858 cells). **(E)** Bar graph showing the cell distribution over the different astrocytic sub-clusters. **(F)** Heatmap showing the differentially genes between the different astrocytic sub-clusters.

To investigate how long-term astrocyte activation alters their gene expression in more detail, we re-clustered the astrocyte cluster (Figure 2D). Sub-clustering of the astrocytes showed that cell types observed in the saccharin group (clusters 0, 2 and 3) were nearly absent from the CNO group, and vice versa (clusters 1 and 4, Figure 2D). This was confirmed by calculating the cell distribution over the different clusters. Cluster 0 accounted for 53.26% of all cells in the saccharin group while only 3.96% in the CNO group. Cluster 1, on the other hand, comprised only 0.87% of cells in the saccharin group, but 66.55% in the CNO group (Figure 2E and Supplementary Table 2). Genes that were differentially expressed between the different astrocytic sub-clusters, as well as gene enrichment and functional annotation analysis of these genes showed the presence of transcriptionally and functionally different astrocytes (Figure 2F). The clusters in the saccharin group (cluster 0, 2 and 3) are enriched for genes involved in regulation of neuronal death, neuroinflammatory response and regulation of synaptic organization, while those of the CNO clusters (cluster 1 and 4) are enriched for genes involved in low-density lipoprotein particles, central nervous system development, gliogenesis, and glial cell differentiation (Supplementary Data 2).

### Chronic activation of astrocytes changes the transcriptome of neighboring microglia

Even though Gq-DREADD was only expressed in astrocytes, we observed a 35% increase and a 42% decrease in the number of cells in the microglia I and II clusters, respectively, after astrocyte activation with CNO (Figure 1E and Supplementary Table 1). In addition, it is known that the increase of intracellular Ca^2+^ after astrocytic activation results in calcium waves that can activate neighboring astrocytes and microglia (Figure 3A). Previous research has shown that activated astrocytes release adenosine triphosphate (ATP) that can activate the purinergic receptors on neighboring microglia, resulting in microglial activation (Figure 3A) (Davalos et al., 2005; Liu et al., 2011; Schipke et al., 2002; Verderio and Matteoli, 2001). As a result, we decided to study the effect of long-term astrocyte activation on the transcriptome of neighboring microglia.

**Figure 3:**
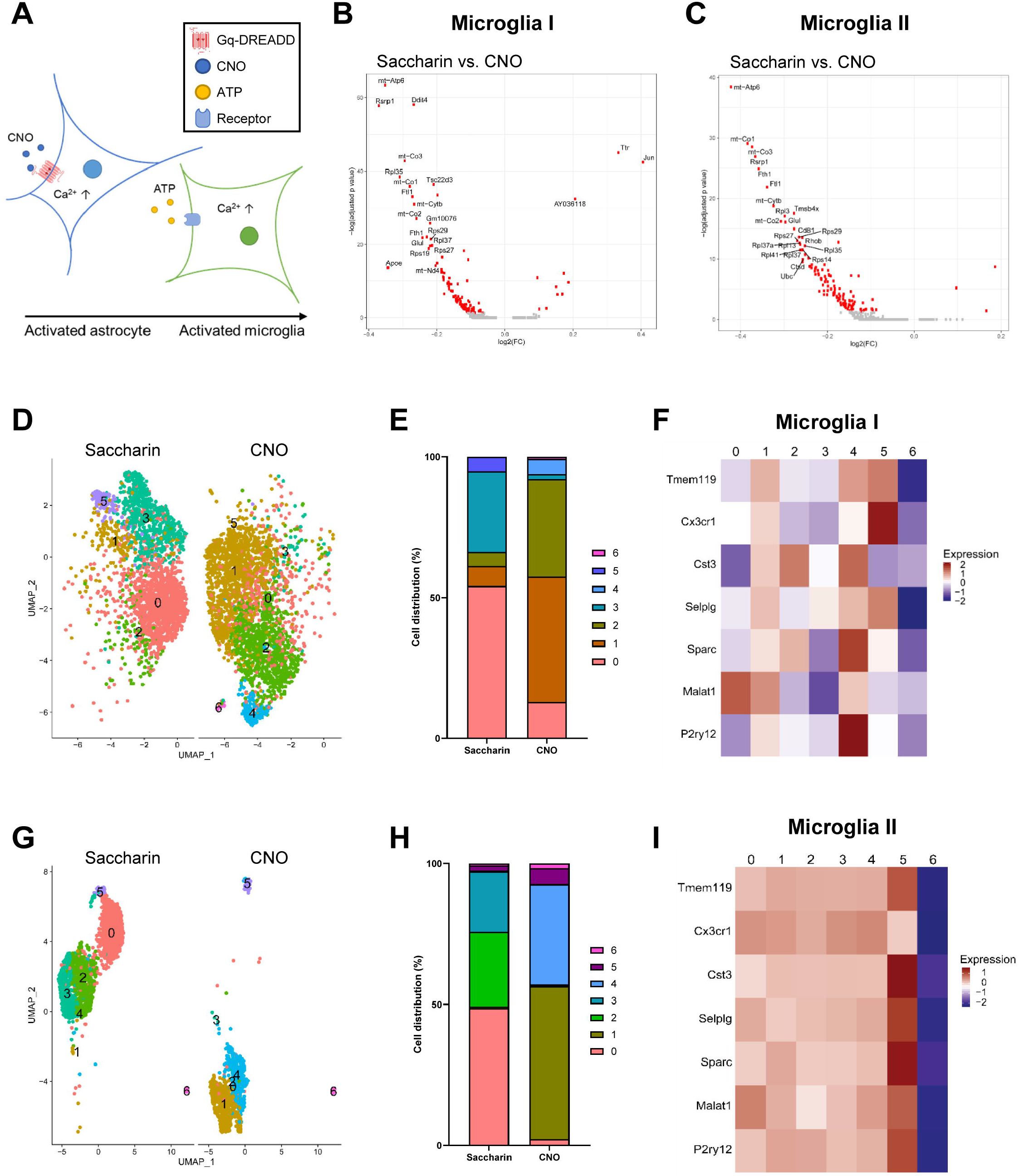
Re-clustering of the microglia I and II cell clusters shows the presence of transcriptionally different microglia after astrocyte-specific activation with the Gq-DREADD. **(A)** Schematic overview showing how activation of astrocytes can lead to activation of neighboring microglia. CNO binds to the Gq-DREADD expressed on astrocytes, leading to an increase of intracellular Ca^2+^. These activated astrocytes release ATP, which activates the purinergic receptors on neighboring microglia. Activation of these purinergic receptors results in an increase of intracellular Ca^2+^ and microglial activation. (Adapted from (Liu et al., 2011)). **(B & C)** Volcano plots showing differentially expressed genes between the saccharin group and the CNO group in the microglia I **(B)** and microglia II **(C)** clusters. Differentially expressed genes with an adjusted p value < 0.05 after Bonferroni correction are shown in red. **(D & G)** UMAP showing the seven clusters identified after re-clustering the microglia I (4,987 cells, **(D)**) and II clusters (3,172 cells, **(G)**). The UMAP is split into two groups, the saccharin group and the CNO group (microglia I (2,293 vs. 2,694 cells **(D)**), microglia II (2,107 cells vs. 1,065 cells **(G)**)). **(E & H)** Bar graph showing the cell distribution over the different microglia I **(E)** and II **(H)** sub-clusters. **(F & I)** Heatmap showing the expression of homeostatic microglial genes in microglia clusters I **(F)** and II **(I)**.

We identified five different microglial clusters (Figure 1D) and noticed that the majority of the differentially expressed genes in the microglial clusters were downregulated (Figure 3B, 3C and Supplementary Figure 3A-C) in the CNO group compared to the saccharin group. Since we observed a 35% increase in microglia I and a 42% decrease in microglia II after CNO treatment, we focused on these microglial clusters first. Gene enrichment and functional analysis of genes that were downregulated after CNO treatment in the microglia I cluster were involved in programmed cell death, and the innate and adaptive immune system, while upregulated genes were involved in the negative regulation of cell growth, cellular metabolic processes, and gene expression (Figure 3B and Supplementary Data 1). Genes that were downregulated after CNO treatment in the microglia II cluster were mainly involved in ribosomal and apoptotic processes while the upregulated genes were involved in MAPK activation and interleukin 17 signaling (Figure 3C and Supplementary Data 1).

Re-clustering of the microglia I cluster resulted in seven sub-clusters of which clusters 4 and 6 were only observed in the CNO group (Figure 3D, 3E and Supplementary Table 3). Differentially expressed genes in these two clusters were involved in G protein-coupled purinergic nucleotide receptor activity, the innate immune system, antigen processing and presentation, and neuronal cell death (Supplementary Data 3). In addition, clusters 0, 3 and 5 were more common in the saccharin group, while clusters 1 and 2 were more pronounced in the CNO group (Figure 3E and Supplementary Table 3). Clusters 1 and 2 showed an enrichment for functional terms involving ribosomal processes, while no gene ontology analysis was performed for clusters 0, 3 and 5 due to the low number of differentially expressed genes (Supplementary Data 3). Gene expression analysis showed a relative upregulation of homeostatic microglial genes in clusters 1, 4 and 5, while they were downregulated in clusters 0, 2, 3 and 6 (Figure 3F).

Re-clustering of the microglia II cell cluster resulted in the identification of seven sub-clusters. Three sub-clusters were mainly observed in the saccharin group (clusters 0, 2 and 3), two were more common in the CNO group (clusters 1 and 4), and two others (clusters 5 and 6) were present in both groups (Figure 3G, 3H and Supplementary Table 3). Differentially expressed genes in clusters 0, 2 and 3 were involved in G protein-coupled purinergic nucleotide receptor activity, the innate immune system and cell death (Supplementary Data 4). Clusters 1 and 4, on the other hand, were enriched for transcripts involved in the innate immune system, lysosomal functions, synaptic pruning, respiratory chain complex, and response to lipoprotein particles (Supplementary Data 4). A strong downregulation of homeostatic microglial genes was observed in cluster 6, while an upregulation was observed in cluster 5 (Figure 3I).

In the microglia III cluster, gene enrichment and functional analysis of the differentially expressed genes showed an enrichment for transcripts involved in the immune system and cell killing, while a downregulation of transcripts involved in monocyte differentiation was observed (Supplementary Figure 3A and Supplementary Data 1).

Re-clustering of the microglia III cluster resulted in five sub-clusters (Figure 4A). Although all of them were observed in both groups, cluster 0 was six times more prominent in the saccharin group (56.08%) compared to the CNO group (8.95%), and cluster 2 was nine times more pronounced in the CNO group (36.24%) than in the saccharin group (3.97%, Figure 4A, 4B and Supplementary Table 3). Gene ontology and functional analysis of the differentially expressed genes pointed towards similar functions for all cells in the different clusters. The differentially expressed genes in clusters 0 and 2 were involved in G protein-coupled receptor binding and the migration of immune cells like neutrophils, granulocytes, and macrophages (Supplementary Data 5). Again, we observed a strong downregulation of homeostatic genes in one cluster, cluster 4, compared to the other clusters (Figure 4E).

**Figure 4:**
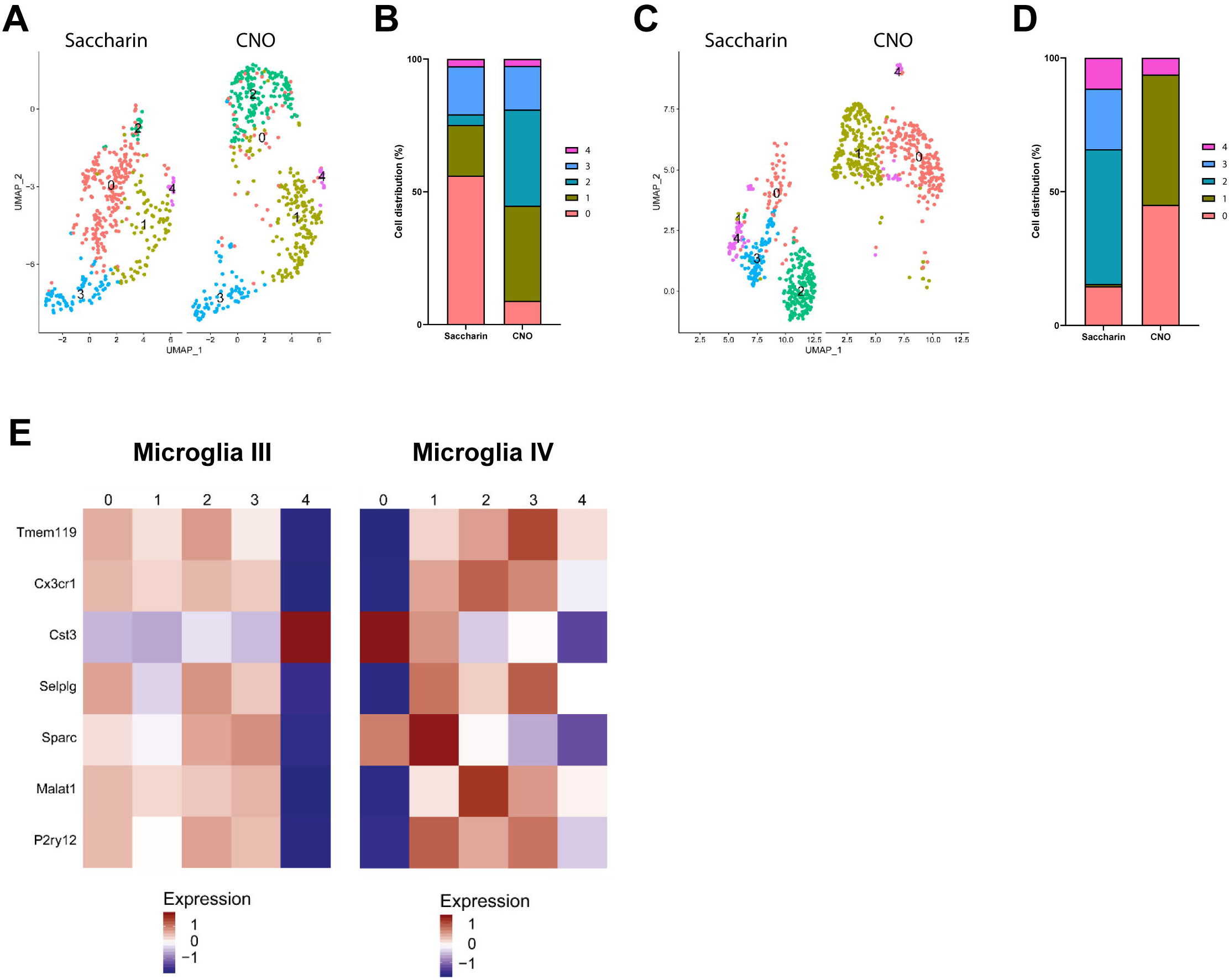
Re-clustering of the microglia III and IV cell clusters shows transcriptionally different cell types after activation of astrocytes with Gq-DREADD. **(A & C)** UMAP showing the different clusters that were identified after re-clustering the microglia III (861 cells**, (A)**) and IV (823 cells **(C)**) clusters. The UMAP is split into two groups, the saccharin group and the CNO group (microglia III (403 vs. 458 cells **(A)**) and IV (375 vs. 448 cells **(C)**). **(B & D)** Bar graph showing the cell distribution over the different microglial sub-clusters (III **(B)** and IV **(D)**). **(E)** Heatmap showing the differential expression of homeostatic microglia genes in microglia clusters III and IV.

Interestingly, the genes that were downregulated in the microglia IV cluster after astrocyte activation were involved in G protein-coupled purinergic nucleotide receptor activity and ATP synthesis coupled electron transport. The genes that were upregulated in the CNO group compared to the saccharin group were enriched for terms involving ribosomal processes (Supplementary Figure 3B and Supplementary Data 1).

The microglia IV cluster showed five sub-clusters after re-clustering (Figure 4C). Clusters 2 and 3 were only observed in the saccharin group, and thus, were completely absent from the CNO group (Figure 4C, 4D and Supplementary Table 3). No gene ontology and functional analysis was performed for the differentially expressed genes in cluster 2 due to the low number of differentially expressed genes. However, the differentially expressed genes in cluster 3 were shown to be involved in gene expression, translation and ribosomal processes (Supplementary Data 6). Cluster 0 (14.67% in the saccharin group vs. 45.09% in the CNO group) and cluster 1 (0.80% in the saccharin group vs. 48.61% in the CNO group) were more common in the CNO group compared to the saccharin group (Figure 4C, 4D and Supplementary Table 3). Differentially expressed genes in these two clusters were involved in abnormal immune system morphology and ubiquitination (Supplementary Data 6). For microglia IV, we observed again that one cluster, cluster 0, showed a downregulation of homeostatic gene expression compared to the other four clusters (Figure 4E).

Lastly, we noticed that genes involved in cell surface receptor signaling pathway and response to lipids were upregulated in the microglia V cluster. The downregulated genes were enriched for gene ontology terms including ribosomal processes (Supplementary Figure 3C and Supplementary Data 1).

The re-clustering of the fifth microglial cluster resulted in four different cell clusters (Supplementary Figure 3D). Clusters 0 and 2, showed a 1.7-fold increase and a 50% decrease in cell numbers between the saccharin and the CNO groups, respectively (Supplementary Figure 3E and Supplementary Table 3). The gene enrichment and functional annotation of the differentially expressed genes for cluster 0 showed that these cells were responsive to cytokines, while cluster 2 showed an involvement in major histocompatibility complex (MHC) binding. In addition, due to the high number of ribosomal genes that were differentially expressed, both clusters also showed an involvement in ribosomal processes (Supplementary Data 7). Same as with microglia I-IV, we noticed that a big difference in expression levels of homeostatic microglial genes. Cluster 0, 2 and 3 showed relatively low expression of homeostatic genes, while cluster 1 showed an increased expression of these genes compared to the other three clusters (Supplementary Figure 2F).

### Re-clustering of the mixed glia cell clusters showed the presence of microglia-like, astrocyte-like and oligodendrocyte-like cells

We annotated two cell clusters as ‘Mixed Glia’ since it was not clear from their conserved marker genes whether these cells should be annotated as astrocytes (e.g. *Cpe* and *Apoe*), microglia (e.g. *Actb*) or oligodendrocytes (e.g. *Dpysl2*; Figure 5A and 5B). Activation of astrocytes using CNO led to a 48% increase in cell numbers in the mixed glia I cluster and a 50% decrease in the mixed glia II cluster (Figure 1E and Supplementary Table 1).

**Figure 5:**
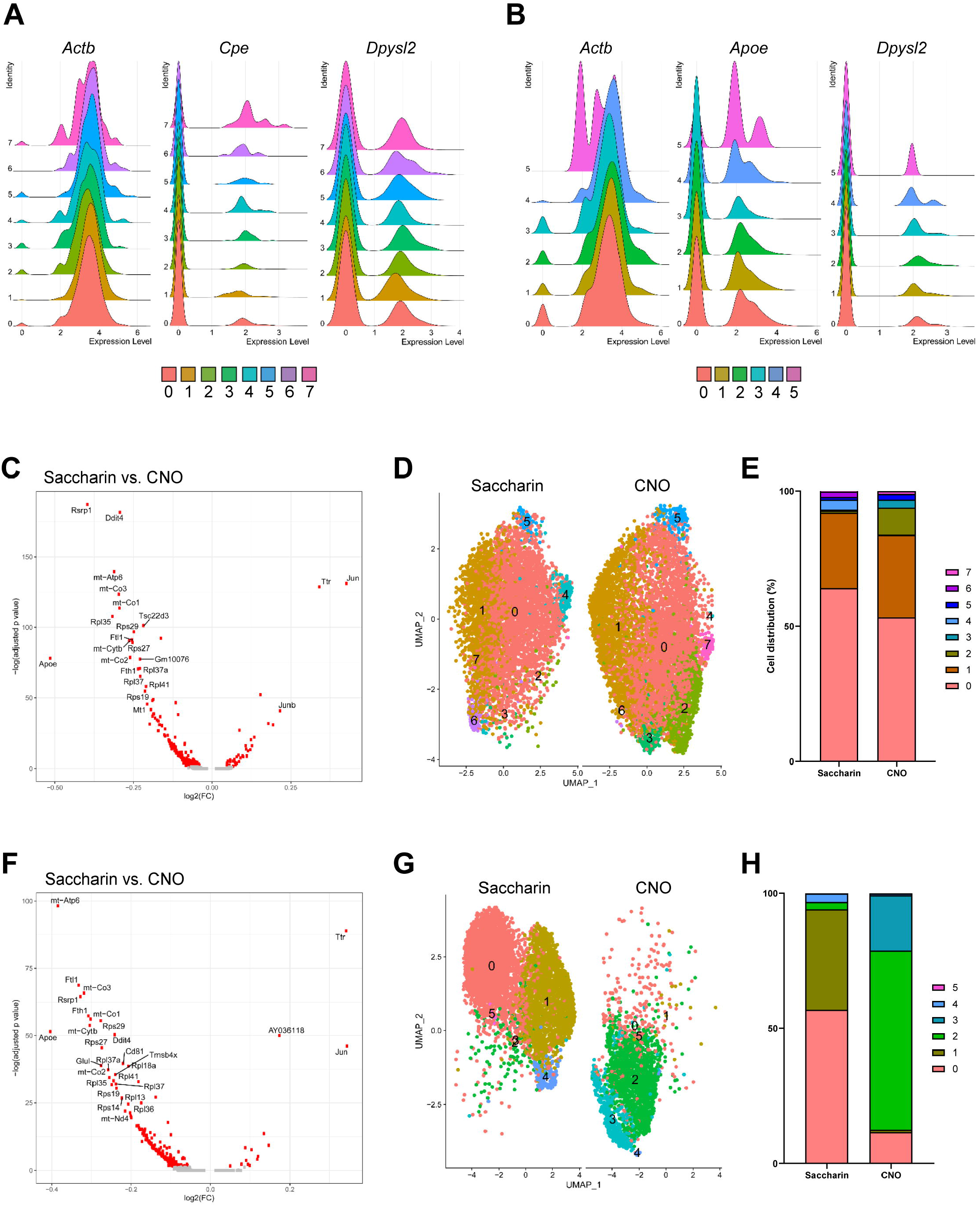
Re-clustering of the mixed glia cell clusters shows transcriptionally different cell types after activation of astrocytes with the Gq-DREADD. **(A & B)** Ridge plots showing the expression of conserved marker genes in the mixed glia I **(A)** and mixed glia II **(B)** clusters. The microglia marker *Actb*, the astrocytic marker *Cpe* and the oligodendrocyte marker *Dpysl2* are expressed in the mixed glia I clusters **(A)**, while the microglial marker *Actb*, the astrocytic marker *Apoe* and the oligodendrocyte marker *Dpysl2* are expressed in the mixed glia II clusters **(B). (C & F)** Volcano plot showing differentially expressed genes between the saccharin group and the CNO group in the mixed glia I **(C)** and II **(F)** clusters. Differentially expressed genes with an adjusted p value < 0.05 after Bonferroni correction are shown in red. **(D)** UMAP showing the eight clusters identified after re-clustering the mixed glia I cluster (13,029 cells). The UMAP is split into two groups, the saccharin group (5,744 cells) and the CNO group (7,285 cells)). **(E & H)** Bar graph showing the cell distribution over the different mixed glial I **(E)** and II **(H)** sub-clusters. **(G)** UMAP showing the six clusters identified after re-clustering the mixed glia II cluster (7,905 cells). The UMAP is split into two groups, the saccharin group (5,504 cells) and the CNO group (2,401 cells).

Differentially expressed genes in the mixed glia I cluster showed that chronic activation of astrocytes by the administration of CNO downregulated genes involved in apoptotic and ATP metabolic processes (Figure 5C and Supplementary Data 1). Genes that were upregulated after CNO treatment are involved in immune cell differentiation and migration, cytokine production, and response to lipids (Figure 5C and Supplementary Data 1). In the mixed glia II cluster, gene enrichment and functional annotation of differentially expressed genes demonstrated that downregulated genes were also involved in apoptotic and ATP metabolic process processes while the genes that were upregulated after Gq-DREADD-mediated activation of astrocytes were mainly involved in gene expression and immune-related processes (Figure 5F and Supplementary Data 1). Based on gene ontology analysis, we expect the presence of both astrocytes and microglia in the mixed glial clusters.

To determine which type of cell might be responsible for the observed effects, we re-clustered both mixed glia clusters. Re-clustering of the mixed glia I cluster resulted in eight different sub-clusters while the re-clustering of the mixed glia II cluster resulted in six sub-clusters (Figure 5D and 5G). Cells belonging to cluster 2 in the mixed glia I group were more abundant in the CNO group compared to the saccharin group. 0.78% of the cells in the saccharin group were part of cluster 2, while 10.06% of cells in the CNO group (Figure 5E and Supplementary Table 4). No gene ontology analysis could be performed for the differentially expressed genes in cluster 2 since only *Apoe* was differentially expressed (Supplementary Data 8). Other clusters that were more pronounced the CNO group compared to the saccharin group are clusters 3 (0.37% in the saccharin group vs. 2.95% in the CNO group), 5 (1.01% in the saccharin group vs. 2.05% in the CNO group) and 7 (0.12% in the saccharin group vs. 1.00% in the CNO group, Supplementary Table 4). Gene ontology analysis showed that these cells are involved in antigen processing and presentation (cluster 3), ribosomal processes (cluster 5), and regulation of transport (cluster 7, Supplementary Data 8).

Re-clustering of the mixed glia II cluster showed more dramatic effects of Gq-DREADD-mediated activation of astrocytes with CNO on the different sub-clusters than was observed in the mixed glia I cluster. Cluster 0 (56.90% in the saccharin group vs. 11.66% in the CNO group), cluster 1 (37.12% in the saccharin group vs. 0.87% in the CNO group) and cluster 4 (3.13% in the saccharin group vs. 0.62% in the CNO group) disappeared almost entirely after CNO treatment while cluster 2 (2.69% in the saccharin group vs. 66.26% in the CNO group) and cluster 3 (0.05% in the saccharin group vs. 20.45% in the CNO group) appeared after CNO treatment (Figure 5G, 5H and Supplementary Table 4). No gene ontology analysis was performed on clusters 1 and 2 since there was only one differentially expressed gene in each cluster (Supplementary Data 9). Differentially expressed genes in clusters 0 and 4 were involved in the response to axon injury and extracellular stimuli, cell motility, and regulation of the actin cytoskeleton (Supplementary Data 9). The genes that were differentially expressed in cluster 3 played a role in ATP metabolic processes, aging and the mitochondrial respiratory complex (Supplementary Data 9).

## Discussion

To the best of our knowledge, this is the first study to show transcriptional changes in glia that occur after long-term activation of astrocytes with the Gq-DREADD in an *in vivo* model. Previous studies have expressed the Gq-DREADD specifically in astrocytes using the GFAP promotor and showed an increase in intracellular Ca^2+^ after activation with CNO (Agulhon et al., 2013; Bull et al., 2014; Scofield et al., 2015). However, none of these studies looked at the effect of long-term Gq-DREADD-mediated activation on the transcriptome of astrocytes or other neighboring brain cells. We performed scRNA seq on hippocampal and cortical cells of C57BL/6J mice to determine the effect of long-term activation of astrocytes through Gq-DREADD. Network analysis of the genes that were upregulated after Gq-DREADD activation with CNO in astrocytes showed an involvement of these genes in the GPCR signaling pathway and calcium homeostasis (Figure 2B and 2C). These findings confirmed the transcriptionally activated state of the astrocytes after long-term activation of Gq-DREADD with CNO. Furthermore, our results demonstrate the presence of transcriptionally different astrocyte populations after Gq-DREADD activation with CNO compared to the control group. Although the up- or downregulation of differentially expressed genes was relatively low, we could clearly observe transcriptionally different astrocytic sub-clusters (Figure 2F). While some clusters were predominantly present in the control group (clusters 0, 2 and 3, Figure 2D and 2E), others were mainly observed after activation with CNO (clusters 1 and 4, Figure 2D and 2E). We also observed that our astrocyte population in the CNO group did not show the transcriptional profile of reactive astrogliosis that is typically seen in neurodegenerative diseases (Supplementary Figure 2D) (Kraft et al., 2013). Furthermore, we also noticed that these Gq-DREADD activated astrocytes did not show the transcriptional profile of other, recently identified astrocyte sub-types such as PAN reactive, A1-specific, A2-specific, disease-associated astrocytes, or astrocytes seen in acute injury and chronic neurodegeneration (Supplementary Figure 2A-C) (Das et al., 2020; Habib et al., 2020; Liddelow et al., 2017). Therefore, we hypothesize that long-term activation of astrocytes changes their transcriptional profile to a, thus far transcriptionally uncharacterized phenotype. Further research is needed to assess what this new transcriptional phenotype means to the astrocytes, their environment, and their biological relevance.

Gene and functional analysis of the astrocyte sub-clusters showed a functional difference between the astrocytes present in the saccharin group and after long-term Gq-DREADD-mediated activation. The astrocyte clusters that were more abundant in the saccharin group than in the CNO group (clusters 0, 2 and 3, Figure 2E) were involved in the regulation of several processes such as transport, neuronal cell death, and synapse structure and activity (Supplementary Data 2). The cell clusters that were more abundant in the CNO group (clusters 1 and 4, Figure 2E) were important for low-density lipoprotein homeostasis and central nervous system development. Strikingly, gene ontology showed that the differentially expressed genes from the CNO sub-clusters were enriched for glial cell development, astrocyte differentiation and positive regulation of neuronal apoptotic processes (Supplementary Data 2). Furthermore, a recent study (Batiuk et al., 2020) determined the transcriptomic profile of astrocytes in the hippocampus and cortex of C57BL/6J mice at post-natal day 56 and showed the existence of five transcriptionally different astrocytic clusters. We looked at the transcriptomic profile of these five astrocyte types in our data and were only able to link our astrocyte sub-cluster 4 to their astrocyte cluster 5 (Supplementary Figure 2E). These astrocytes seem to be enriched in cortical layers 2/3 and 5, and in the dentate gyrus of the hippocampus (Batiuk et al., 2020). While the astrocytes enriched in the saccharin group show a homeostatic phenotype (Barres, 2008), the phenotype of the astrocytes enriched in the CNO group is less clear. Previous research has shown that the transcriptomic profile of astrocytes varies in response to different “insults”, e.g. long-term activation (Anderson et al., 2014; Hamby et al., 2012). Therefore, additional research is needed to elucidate the exact molecular phenotype of the Gq-DREADD activated-astrocytes.

Furthermore, the transcriptome of neighboring microglia was also dramatically affected by the long-term activation of astrocytes (Figures 3 and 4). We observed microglial clusters that were absent in the control group and dominating in the CNO group (Figures 3D, 3G, 4A and 4C). We also observed that the transcriptomic changes that occurred in microglia after astrocytic Gq-DREADD-mediated activation were different than those in control microglia (Supplementary Data 3-7). After long-term activation of astrocytes, the microglial transcriptome suggests a role in lipoprotein particle processes, and migration and chemotaxis of immune cells. We also observed the presence of microglial sub-clusters enriched for genes important for the purinergic receptor pathway on microglia. As mentioned earlier, activated astrocytes release ATP which activates the purinergic receptors on microglia, resulting in microglial activation (Figure 3A) (Davalos et al., 2005; Liu et al., 2011; Schipke et al., 2002; Verderio and Matteoli, 2001).

Additionally, we observed that certain microglial sub-clusters showed an increased expression of genes that are known to characterize homeostatic microglia, while other sub-clusters showed a decreased expression of these genes (Figure 3F, 3I, 4E and Supplementary Figure 3F). To determine the transcriptomic profile of these microglial sub-cluster, we determined whether these clusters showed the expression profile of disease-associated microglia (DAM). This subgroup of microglia were identified while studying the immune cells at the single cell level in a mouse model for AD (Deczkowska et al., 2018; Keren-Shaul et al., 2017). We observed an upregulation of DAM genes in some of our microglial sub-clusters (Supplementary Figure 4), resulting in the hypothesis that DAM cells may not always be linked to amyloid pathology and thus are not always disease-associated. Our data hypothesizes that DAM genes are induced following astrocyte activation. This would implicate that the microglial response is secondary to the astrocytic response in AD, which can have major implications for AD research. However, further research is needed to elucidate what causes the induction of DAM genes after long-term activation of astrocytes in a healthy mouse model.

We further observed a 48% increase and 50% decrease in cell numbers in the two largest cell clusters, mixed glia I and mixed glia II respectively, after Gq-DREADD-mediated activation of astrocytes. All cell clusters were annotated manually based on marker genes that were conserved between the saccharin group and the CNO group and gene lists present in literature (Arneson et al., 2018; McKenzie et al., 2018). Still, about 60% of the cells in our analysis were annotated as “Mixed Glia”. A possible explanation for this phenomenon is that we analyzed the cortex and hippocampus together, while it is well established that both astrocytes and microglia show regional heterogeneity at the transcriptome level (Batiuk et al., 2020; Tan et al., 2020). Gene ontology analysis of differentially expressed genes in both mixed glial clusters pointed towards the presence of mainly microglia. The dominance of microglia genes was confirmed when we looked at the expression of cell-type-specific marker genes for microglia (e.g. *Cx3cr1*, *P2ry12*, *Tmem119*, *Aif1*, *Olfml3*, *Ccl3*, *Itgam*), astrocytes (e.g. *Aldh1l1*, *Atp1b2*, *Aqp4*, *Sox9*, *Slc4a4*, *Mlc1*) and oligodendrocytes (e.g., *Nfasc*, *Kndc1*; Supplementary Figure 5) (Pan et al., 2020). These difficulties in annotating cell clusters in brain shows the urgency for reliable automatic annotation tools.

Previous research also showed that activation of astrocytes in the mouse hippocampus using the Gq-DREADD influences the excitability of neurons. The increased intracellular Ca^2+^ levels stimulated glutamate release which resulted in activation of neuronal NMDA receptors (Durkee et al., 2019). Since less than 1% of our total cell population subsists of neuronal cells, we were unable to confirm this neuronal activation at the transcriptomic level. Additional research with a different cell isolation protocol is needed to study the effect of long-term activation of astrocytes on the transcriptomic profile of neurons.

Lastly, our single cell dissociation protocol shows a dramatic enrichment of glial cells and a relative depletion of neurons. Possible explanations for this glial enrichment are (1) the fact that we dissociated brain tissue of four months-old mice. Previous studies mostly perform scRNA seq in younger mice, while single nucleus RNA sequencing is more common in adult mice (Hook et al., 2018; Loo et al., 2019; Zeisel et al., 2015; Zhou et al., 2020). The central nervous system of older mice is more complex than that of younger mice, making it harder to dissociate it into single cells, and as a result some cell types can get over- or underrepresented (Darmanis et al., 2015; Grindberg et al., 2013; Krishnaswami et al., 2016; Lake et al., 2016; Lake et al., 2017; Tasic et al., 2018). (2) To arrest ongoing gene expression and to decrease the likelihood of inducing heat shock proteins or immediate-early response genes, we performed our dissociation protocol at 4°C instead of 37°C. Research has also shown that the enzymatic dissociation of brain tissue at 37°C results in an upregulation of inflammatory genes in microglia. The phenomenon was not observed with a mechanical dissociation at 4°C (Bennett et al., 2016). However, one drawback of an ice-cold dissociation is the decrease in efficiency of dissociation with certain tissues such as the brain (Adam et al., 2017; Denisenko et al., 2019; van den Brink et al., 2017). As such, one might hypothesize that ice-cold dissociation was not capable of breaking neuronal connections, while the connections formed by glial cells were easier to break.

In summary, our data show for the first time the effect of long-term activation of astrocytes with Gq-DREADD. We have shown that prolonged activation of astrocytes alter the transcriptional profile of astrocytes and neighboring microglia in the adult mouse brain. Our findings are also important for the interpretation of future studies using Gq-DREADD activation in astrocytes. Finally, our data provides an important resource for future chemogenetic studies and demonstrates that studies of cell-type conditional DREADDs expression would do well to consider other cell types within the tissue of interest.

## Acknowledgments

We thank Dr. John Fryer and Jonathon Sens for the single-cell isolation protocol and Dr. Leonard Petrucelli, Karen Jansen-West, and Lillian Daughrity for virus packaging of the plasmid (Department of Neuroscience, Mayo Clinic). We also thank Dr. Luke C. Dabin, Dr. Md Mamun Al-Amin and Dom J. Acri for their input on the data. This publication was made possible by the Stark Neurosciences Research Institute, the Indiana Alzheimer Disease Center, Eli Lilly and Company, and by the Indiana Clinical and Translational Sciences Institute, funded in part by grant # UL1TR002529 from the NIH, National Center for Advancing Translational Sciences (awarded to S.P.). Sequencing was carried out in the Center for Medical Genomics at Indiana University School of Medicine, which is partially supported by the Indiana Genomic Initiative at Indiana University (INGEN); INGEN is supported in part by the Lilly Endowment, Inc. The authors also acknowledge the Indiana University Pervasive Technology Institute for providing Carbonate supercomputer resources (Stewart et al., 217). This work was supported, in part by grants from Strategic Research Initiative, Precision Health Initiative (Indiana University), NIH R01AG054102, R01AG053500, R01AG053242, and R21AG050804, the National Science Foundation Grant No. CNS-0521433, and Shared University Research grants from IBM, Inc., to Indiana University. The content is solely the responsibility of the authors and does not necessarily represent the official views of the National Institutes of Health, National Science Foundation or Eli Lilly and Company. Figure 1A uses an image from Servier Medical Art, Servier Co.

## Author Contribution

Conceptualization, S.P, M.T.T. and J.K.; Methodology, M.T.T. and J.K.; Formal Analysis, S.P.; Investigation, S.P., M.T.T., B.P.T. and Y.M.; Data Curation, S.P.; Writing – Original Draft, S.P. and M.T.T.; Writing – Review & Editing, S.P., M.T.T., Y.M. and J.K.; Visualization, S.P.; Funding Acquisition, S.P. and J.K.; Supervision, J.K.

