## Supplementary file for "Single-cell resolution analysis of the crosstalk between chemogenically activated astrocytes and microglia"

**Key words**

### ***Supplementary Figure***

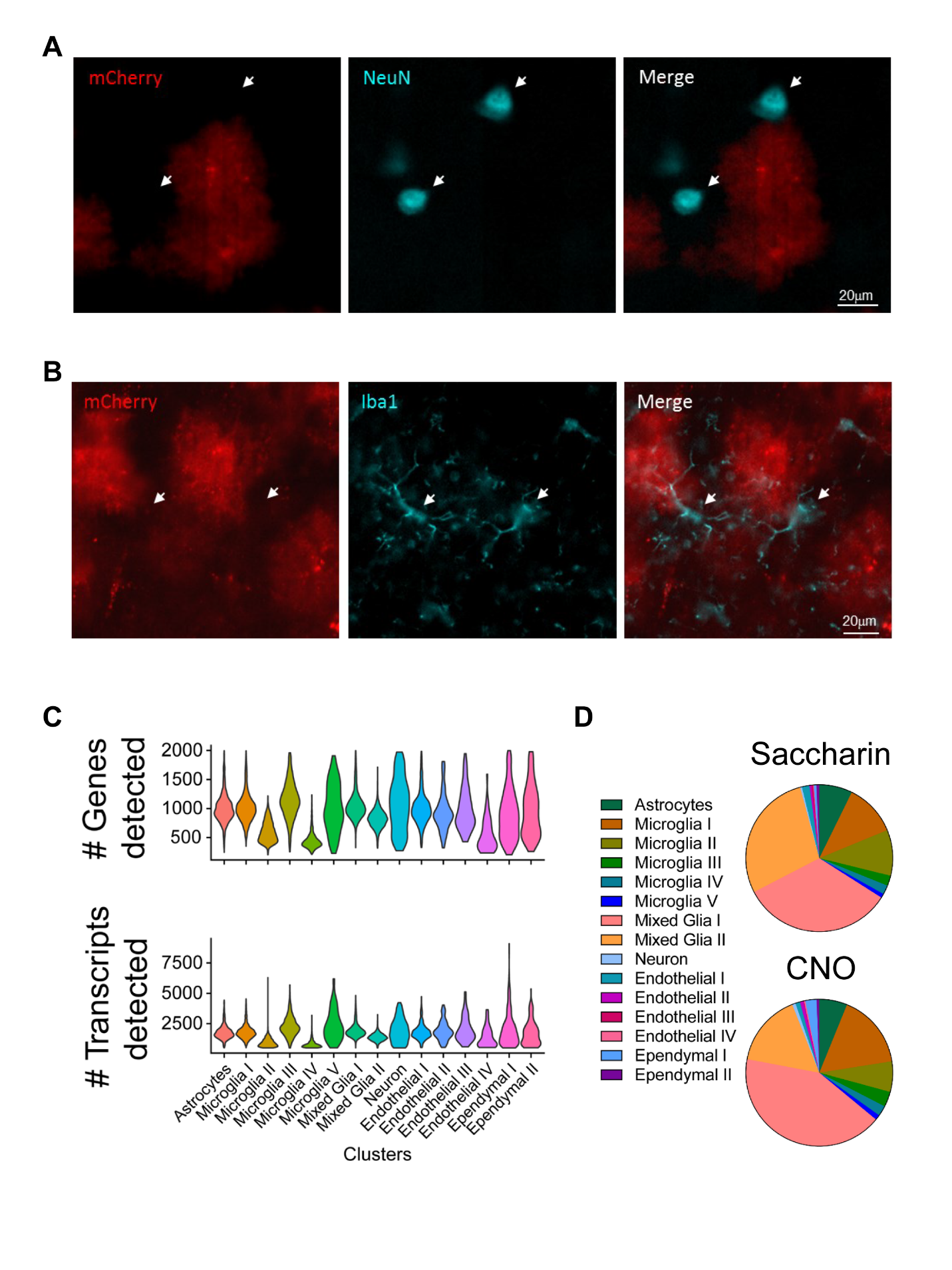

**Supplementary Figure 1: Immunofluorescence showing that Gq-DREADD is not expressed in neurons and microglia and quality control of the single cell RNA sequencing data. (A)** Representative fluorescence images of the hippocampus and cortex demonstrating Gq-DREADD virus spread (red, mCherry) and neurons (cyan, NeuN). **(B)** Representative fluorescence images of the hippocampus and cortex demonstrating Gq-DREADD virus spread (red, mCherry) and microglia (cyan, Iba1). **(C)** Violin plots showing the number of detected genes (top) and transcripts (bottom) in the different cell clusters. **(D)** Pie charts showing the distribution of cells across the different clusters separated per condition (n=2 mice/condition).

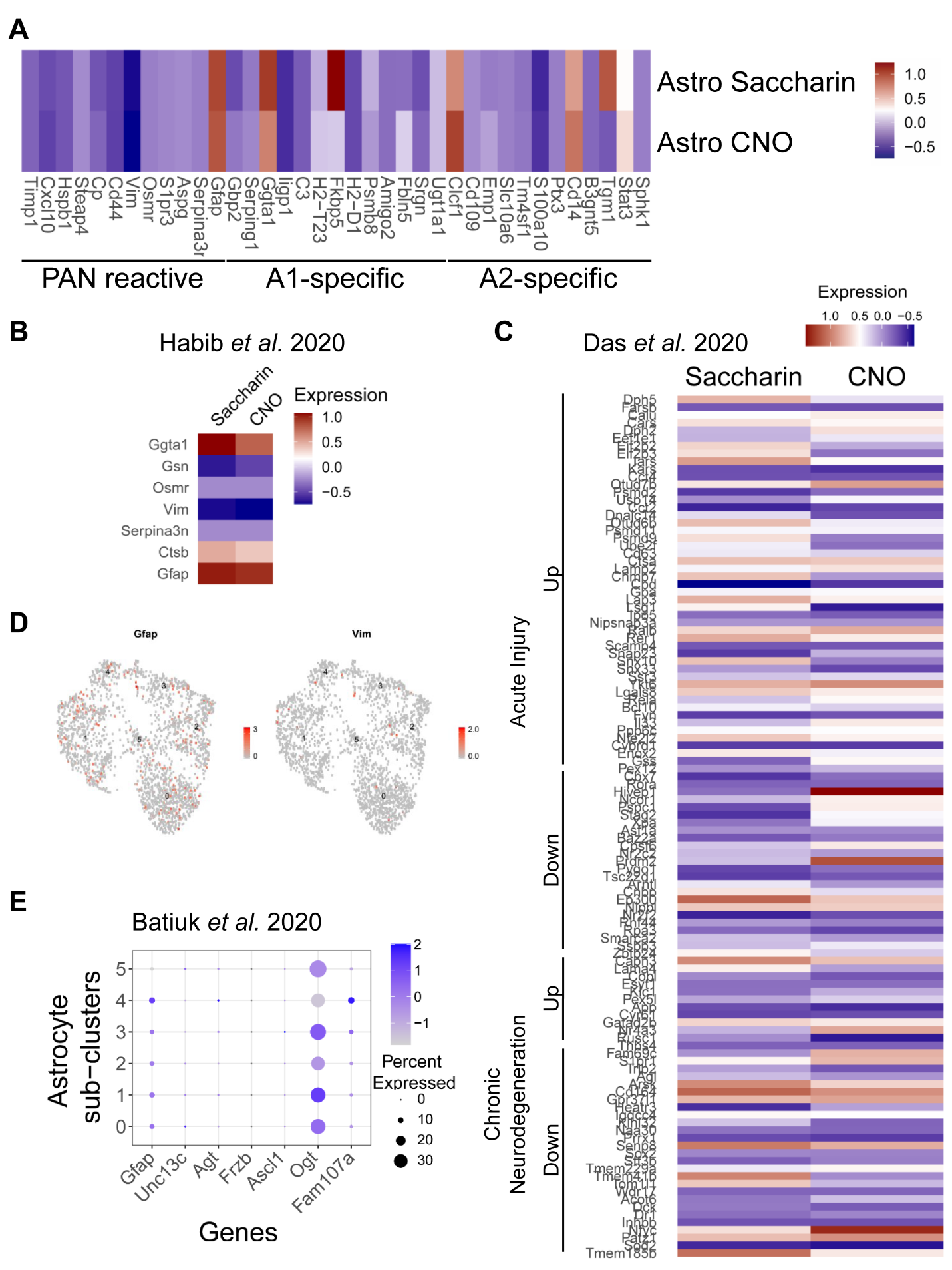

**Supplementary Figure 2: The transcriptomic profile of Gq-DREADD-activated astrocytes are different from previously published profiles. (A)** No upregulation of genes linked to PAN-reactive, A1-specific or A2-specific astrocytes was identified in the Gq-DREADD-activated astrocytes. Marker genes were derived from (Liddelow et al., 2017). **(B & C)** Heatmaps showing the signature genes for disease-associated astrocytes **(B)** and genes that are up- or downregulated in either acute injury or chronic neurodegeneration **(C)** (Das et al., 2020; Habib et al., 2020). Gq-DREADD-activated astrocytes are not enriched for genes involved in any of the phenotypes. **(D)** UMAP of the astrocyte sub-clusters shows very low or no expression of the reactive astrogliosis markers Gfap and Vim. **(E)** Marker genes for the astrocytic clusters identified by Batiuk *et al.* 2020 in the hippocampus and cortex of C57BL/6J (Batiuk et al., 2020).

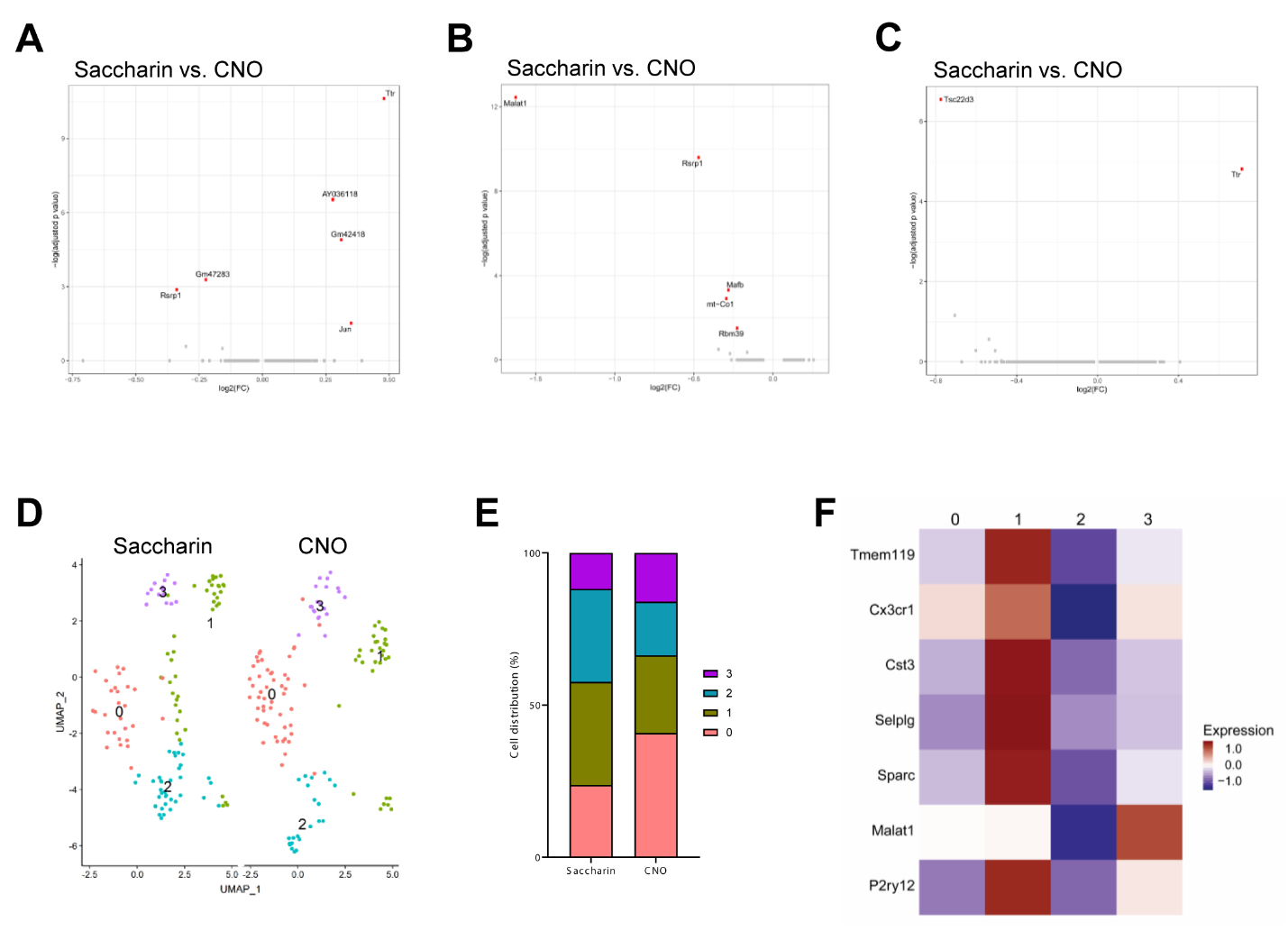

**Supplementary Figure 3: Gq DREADD-mediated activation of astrocytes changes the transcriptome of microglia. (A, B & C)** Volcano plots showing differentially expressed genes between the saccharin group and the CNO group in the microglia III **(A)**, microglia IV **(B)**, microglia V **(C)** clusters. Differentially expressed genes with an adjusted p value < 0.05 after Bonferroni correction are shown in red. **(D)** UMAP showing the different clusters that were identified after re-clustering the microglia V (243 cells) cluster. The UMAP is split into two groups, the saccharin group and the CNO group (118 vs. 125 cells, respectively). **(E)** Bar graph showing the cell distribution over the different microglia V sub-clusters. **(F)** Heatmap showing the expression profile of homeostatic microglial genes in the sub-clusters of microglia V.

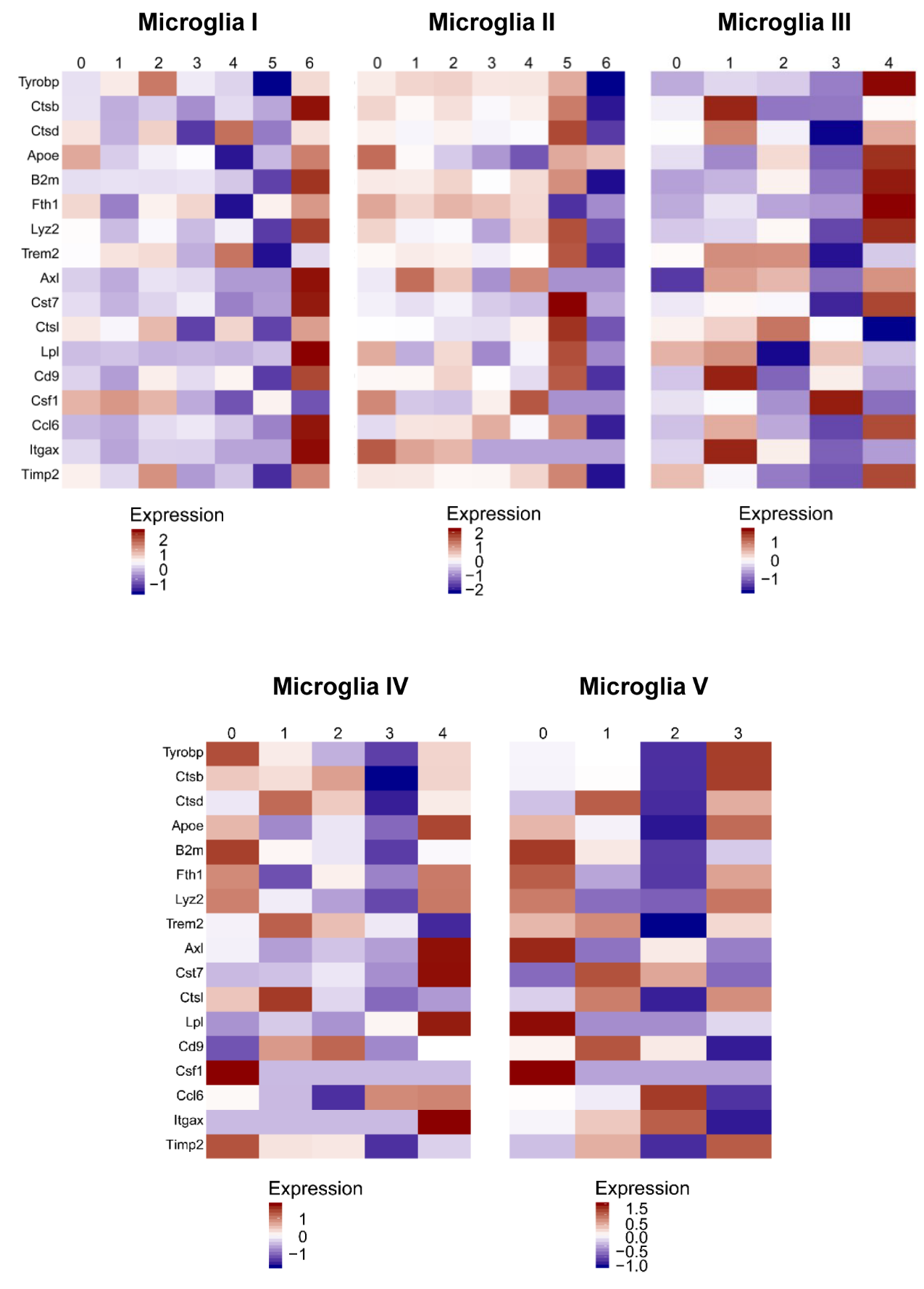

**Supplementary Figure 4: Expression of DAM marker genes in the microglial sub-clusters.** Heatmaps showing the expression profile of DAM genes in the microglial sub-clusters. Microglia I sub-cluster 6, microglia II sub-cluster 5, microglia III sub-cluster 4, microglia IV sub-clusters 0 and 4, and microglia V sub-clusters 0 and 3 showed an increased expression of DAM genes.

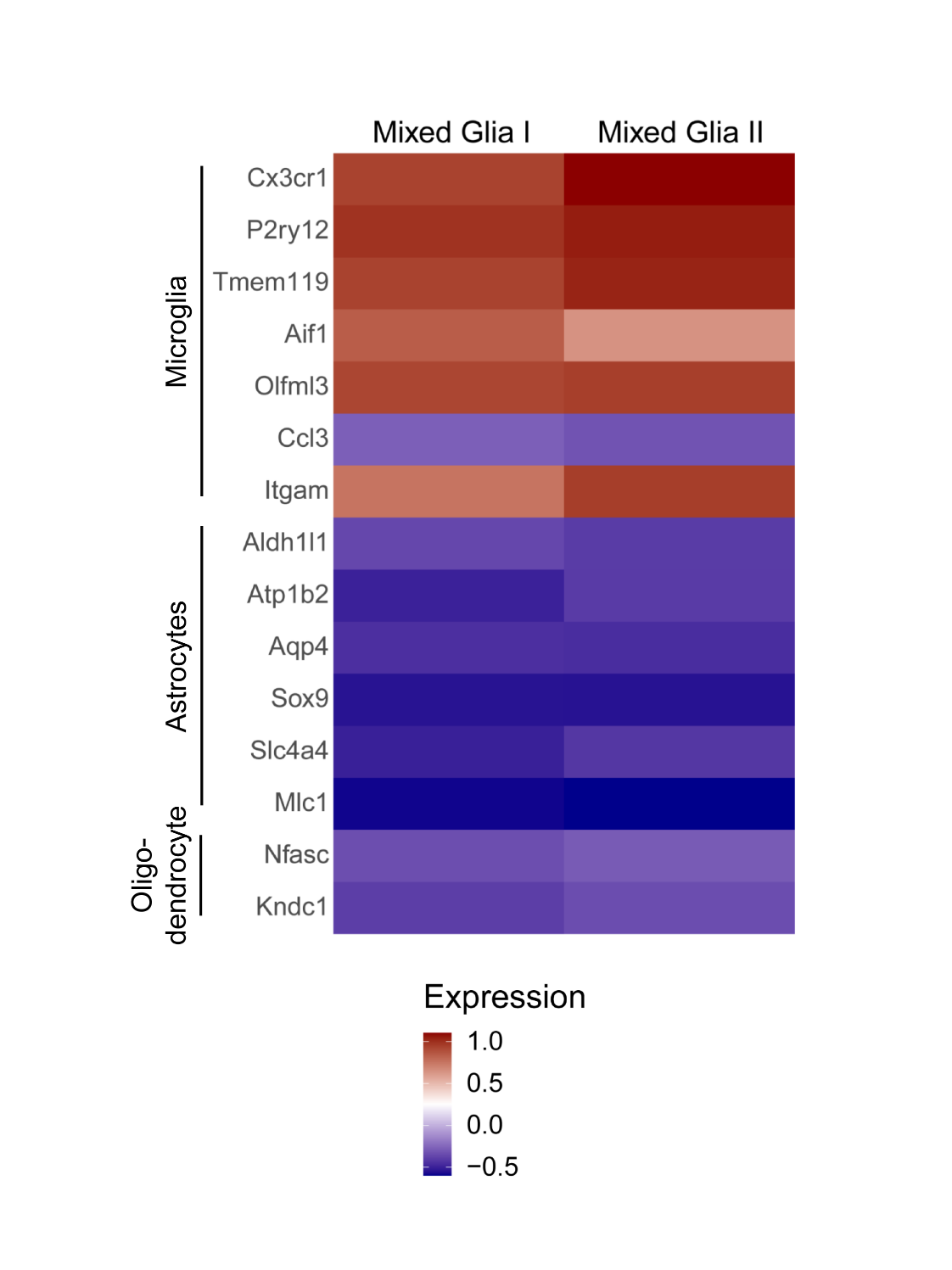

**Supplementary Figure 5: Cell-type-specific marker genes show an upregulation of microglial genes in the mixed glial clusters.** Heatmap showing the expression of cell-type specific markers for microglia, astrocytes and oligodendrocytes in the mixed glia I and mixed glia II clusters.

### ***Supplementary Tables***

Supplementary Table 1: Percentage of cells present in the identified cell clusters

| **Cluster** | **Saccharin (%)** | **CNO (%)** | **Cell Type** |
| --- | --- | --- | --- |
| 0 | 30.66 | 45.32 | Mixed Glia I |
| 1 | 29.62 | 14.87 | Mixed Glia II |
| 2 | 12.37 | 16.74 | Microglia I |
| 3 | 11.39 | 6.60 | Microglia II |
| 4 | 7.41 | 5.30 | Astrocytes |
| 5 | 2.15 | 2.82 | Microglia III |
| 6 | 2.01 | 2.79 | Microglia IV |
| 7 | 1.56 | 1.07 | Endothelial I |
| 8 | 0.43 | 1.64 | Ependymal I |
| 9 | 0.61 | 0.76 | Microglia V |
| 10 | 0.59 | 0.81 | Neuron |
| 11 | 0.46 | 0.43 | Ependymal II |
| 12 | 0.29 | 0.49 | Endothelial II |
| 13 | 0.29 | 0.14 | Endothelial III |
| 14 | 0.11 | 0.17 | Endothelial IV |

Supplementary Table 2: Percentage of cells present in the astrocyte sub-clusters

| **Cluster** | **Saccharin (%)** | **CNO (%)** |
| --- | --- | --- |
| 0 | 53.26 | 3.96 |
| 1 | 0.87 | 66.55 |
| 2 | 26.96 | 1.52 |
| 3 | 14.13 | 0.47 |
| 4 | 1.01 | 18.30 |
| 5 | 3.77 | 9.21 |

Supplementary Table 3: Percentage of cells present in the microglial sub-clusters

| **Cluster** | **Saccharin (%)** | **CNO (%)** |
| --- | --- | --- |
| ***Microglia I*** | | |
| 0 | 54.12 | 12.81 |
| 1 | 7.11 | 44.62 |
| 2 | 5.02 | 38.74 |
| 3 | 28.74 | 1.78 |
| 4 | 0.00 | 5.27 |
| 5 | 5.02 | 0.07 |
| 6 | 0.00 | 0.71 |
| ***Microglia II*** | | |
| 0 | 48.74 | 2.25 |
| 1 | 0.52 | 54.18 |
| 2 | 26.48 | 0.28 |
| 3 | 21.40 | 0.38 |
| 4 | 0.33 | 35.59 |
| 5 | 1.80 | 5.63 |
| 6 | 0.71 | 1.69 |
| ***Microglia III*** | | |
| 0 | 56.08 | 8.95 |
| 1 | 19.11 | 35.81 |
| 2 | 3.97 | 36.24 |
| 3 | 18.11 | 16.38 |
| 4 | 2.73 | 2.62 |
| ***Microglia IV*** | | |
| 0 | 14.67 | 45.09 |
| 1 | 0.80 | 48.66 |
| 2 | 50.40 | 0.00 |
| 3 | 22.67 | 0.00 |
| 4 | 11.47 | 6.25 |
| ***Microglia V*** | | |
| 0 | 23.73 | 40.80 |
| 1 | 33.90 | 25.60 |
| 2 | 30.51 | 17.60 |
| 3 | 11.86 | 16.00 |

Supplementary Table 4: Percentage of cells present in the mixed glia I and II sub-clusters

| **Cluster** | **Saccharin (%)** | **CNO (%)** |
| --- | --- | --- |
| ***Mixed Glia I*** | | |
| 0 | 64.05 | 53.32 |
| 1 | 27.92 | 30.50 |
| 2 | 0.78 | 10.06 |
| 3 | 0.37 | 2.95 |
| 4 | 3.71 | 0.04 |
| 5 | 1.01 | 2.05 |
| 6 | 2.04 | 0.08 |
| 7 | 0.12 | 1.00 |
| ***Mixed Glia II*** | | |
| 0 | 56.90 | 11.66 |
| 1 | 37.12 | 0.87 |
| 2 | 2.69 | 66.26 |
| 3 | 0.05 | 20.45 |
| 4 | 3.13 | 0.62 |
| 5 | 0.11 | 0.12 |

Batiuk, M.Y., Martirosyan, A., Wahis, J., de Vin, F., Marneffe, C., Kusserow, C., Koeppen, J., Viana, J.F., Oliveira, J.F., Voet, T.*, et al.* (2020). Identification of region-specific astrocyte subtypes at single cell resolution. Nature Communications 11, 1220.

Das, S., Li, Z., Noori, A., Hyman, B.T., and Serrano-Pozo, A. (2020). Meta-analysis of mouse transcriptomic studies supports a context-dependent astrocyte reaction in acute CNS injury versus neurodegeneration. Journal of Neuroinflammation 17, 227.

Habib, N., McCabe, C., Medina, S., Varshavsky, M., Kitsberg, D., Dvir-Szternfeld, R., Green, G., Dionne, D., Nguyen, L., Marshall, J.L.*, et al.* (2020). Disease-associated astrocytes in Alzheimer’s disease and aging. Nature Neuroscience.

Liddelow, S.A., Guttenplan, K.A., Clarke, L.E., Bennett, F.C., Bohlen, C.J., Schirmer, L., Bennett, M.L., Munch, A.E., Chung, W.S., Peterson, T.C.*, et al.* (2017). Neurotoxic reactive astrocytes are induced by activated microglia. Nature 541, 481-487.
